# MLL3 adaptor function, not methyltransferase catalytic activity, is essential for breast tumor suppression

**DOI:** 10.64898/2026.06.03.729916

**Authors:** Kenta Nishitani, Jihong Cui, Marcelo Coutinho de Miranda, Guojia Xie, Nicole Couturier, Yuta Matsuno, Masako Suzuki, Gregoire Lauvau, Kai Ge, Wenjun Guo

**Author notes:** **Correspondence to WG:**.

## Abstract

MLL3 (Mixed-Lineage Leukemia 3), also known as KMT2C, is one of the most frequently altered epigenetic regulators in breast cancer. MLL3 loss-of-function leads to accelerated tumor onset and growth and increased metastasis. As a large multi-domain protein, MLL3 functions as a histone methyltransferase and a nuclear protein adaptor interacting with other epigenetic proteins. Since breast cancer MLL3 mutations are often truncating mutations that lead to protein degradation, whether the MLL3 tumor suppressor activity depends on its catalytic activity or non-catalytic chromatin adaptor function remains unclear. Here, using CRISPR genetically engineered mouse mammary stem cell organoid-based breast tumor models, we dissected dosage-dependent and domain-specific functions of MLL3 in breast tumor suppression. MLL3 heterozygous loss breast tumor models revealed that MLL3 is haplo-insufficient for breast tumor suppression. Interestingly, homozygous catalytic-dead MLL3-Y4792A mutation did not accelerate tumor onset, growth, or metastasis. By contrast, G367V mutation in the PHD2 domain, which disrupts the BAP1 complex binding without affecting MLL3 protein stability, accelerated tumor onset and growth, phenocopying MLL3 loss. Mechanistically, MLL3 loss impaired chromatin localization of UTX, and genetic depletion of UTX accelerated breast tumor progression in MLL3-wildtype but not MLL3-deficient cells. Integrated RNA-seq, CUT&TAG, and ATAC-seq analyses further showed that transcriptional changes induced by MLL3 loss were more closely associated with promoter-proximal alterations in H3K27Ac, H3K27me3, and chromatin accessibility than with putative MLL3-dependent enhancer regions. Together, these findings reveal that MLL3 suppresses breast tumor initiation through a dosage-sensitive, catalytic-independent adaptor function that regulates promoter-proximal epigenetic states.

## Introduction

MLL3 (Mixed-Lineage Leukemia 3), also known as KMT2C, is one of the most frequently mutated epigenetic regulator genes in human cancers, including breast cancer^[1, 2]^. The majority of MLL3/KMT2C mutations are loss-of-function alterations, such as truncating mutations and deletions, supporting its role as a tumor suppressor. Clinically, MLL3/KMT2C alterations have been associated with poor outcomes in human cancer, including breast cancer^[3, 4]^ and prostate cancer^[5]^. Consistent with these human cancer associations, mouse genetic studies have demonstrated that loss of MLL3 promotes tumor progression in multiple cancer types, including acute myeloid leukemia, hepatocellular carcinoma, urothelial cancer, and breast cancer^[3, 6–8]^. Recent work has also demonstrated that MLL3 loss accelerates breast tumor formation by establishing an immunosuppressive tumor immune microenvironment through the recruitment and induction of highly differentiated effector regulatory T cells^[9]^. An immunosuppressive function of MLL3 loss has also been shown in squamous cell carcinoma^[10]^. However, how MLL3 suppresses breast tumor formation as a chromatin regulator remains poorly understood.

MLL3 has been classically characterized as a histone methyltransferase that catalyzes mono-(me1) and di-methylation (me2) of histone H3 lysine 4 (K4). Since H3K4me1 is broadly enriched at enhancer regions, MLL3 has been recognized as an enhancer regulator that controls enhancer-dependent transcriptional programs^[1, 11]^. Importantly, MLL3 also functions as a scaffold within chromatin regulatory complexes known as a COMPASS-like complex. Through its interaction with multiple epigenetic regulatory proteins, including the histone acetyltransferase p300 and the H3K27me3 demethylase UTX, MLL3 could mediate global epigenetic landscape beyond its own catalytic activity^[1, 11]^. Of note, recent studies further identified non-canonical binding partners of MLL3, including BAP1^[12]^ and TET3^[13]^, which could directly bind with MLL3 and possess epigenetic regulatory functions, highlighting the non-enzymatic chromatin regulatory functions of MLL3. Studies in embryonic stem cells showed that MLL3 methyltransferase and adaptor functions have distinct roles in gene expression regulation^[14]^.

A major unresolved question is whether the tumor suppressor function of MLL3 depends primarily on its catalytic activity or on its non-catalytic scaffold/adaptor functions. Previous genetic studies, including ours, have largely relied on complete loss of MLL3, which removes the entire protein and abolishes both H3K4 methyltransferase activity and its nuclear adaptor functions. Therefore, these models cannot determine whether MLL3 suppresses tumor progression through its enzymatic regulation of enhancer methylation, through non-catalytic adaptor function for chromatin regulatory complexes, or through both mechanisms. Dissecting MLL3’s domain-specific functions is therefore essential for defining the molecular basis of its tumor suppressor activity.

Here, we used CRISPR genetically engineered mouse mammary stem cell organoid-based breast tumor models^[9, 15]^, which enable rapid generation of breast tumors carrying defined patient-relevant genetic alterations, to dissect the domain-specific functions of MLL3 in breast tumor suppression. We generated breast tumor models harboring either a catalytic-dead mutation or an adaptor site mutation in MLL3 and directly compared their tumorigenic potential with that of MLL3 loss. We found that the catalytic activity of MLL3 is dispensable for breast tumor suppression, whereas disruption of the BAP1-binding site by a G367V mutation phenocopies complete MLL3 loss. Integrated epigenomic and transcriptomic analyses further revealed that MLL3 loss does not primarily alter gene expression through putative MLL3-mediated enhancer regions. Instead, transcriptional changes are more closely associated with altered epigenetic states near transcription start sites, and these promoter-associated changes are recapitulated by the adaptor site mutation. Together, these findings redefine MLL3 as a non-catalytic chromatin adaptor tumor suppressor in breast cancer and identify its BAP1-binding function as a critical mechanism to suppress breast tumor progression.

## Results

### MLL3 is haplo-insufficient for suppressing breast tumor initiation

We first examined the association between *KMT2C*/*MLL3* gene mutation status and breast cancer prognosis using the TCGA-BRCA data set. As expected, the tumors with shallow deletion or point mutation (het loss like) and deep deletion (hom deletion) in *KMT2C* gene showed lower *KMT2C* mRNA expression, with severe reduction by deep deletion (**Fig. 1A**). The patients with hom deletion also had shorter disease-specific overall survival (HR = 4.065), and the het loss like patients also had a trend shorter survival (HR = 1.317) (**Fig. 1B**), suggesting haplo-insufficiency of KMT2C in suppressing breast tumor progression.

**Figure 1.**
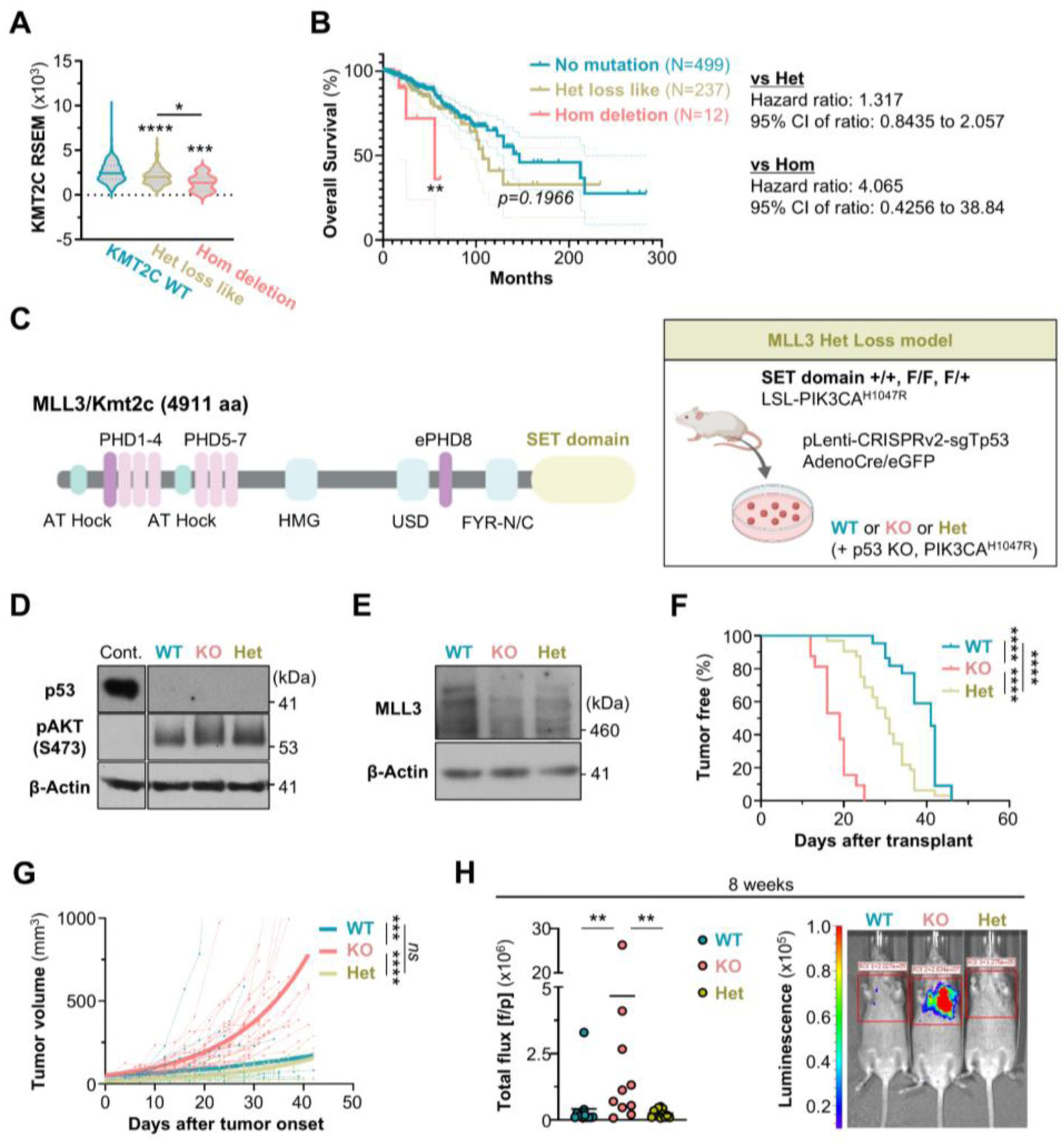
MLL3 is haplo-insufficient for breast tumor initiation, but not for tumor growth and metastasis. **(A)** KMT2C mRNA expression in breast cancer patients (TCGA-BRCA PanCancerAtlas, 1084 patients), stratified by KMT2C mutation status. KMT2C WT, N=324; Het loss like, N=237; Hom deletion, N=12. RSEM, RNA-Seq by Expectation-Maximization. **(B)** Kaplan-Meier overall survival curves of breast cancer patients from the TCGA-BRCA dataset based on KMT2C mutation status. No mutation, N=499; Het loss like, N=237; Hom deletion, N=12. **(C)** Schematic overview of the generation of the MLL3 heterozygous (Het) and homozygous (Hom) loss breast tumor models. **(D)** Western blot (WB) analysis of Trp53 and pAKT (S473). β-Actin was used as an internal control. **(E)** WB analysis of MLL3 protein levels in MLL3 WT, Hom, and Het MaSCs. **(F)** Tumor onset curve with MLL3 WT, Hom, or Het breast tumor models. Three biologically independent experiments were combined. WT, N=22; Hom, N=32; Het, N=32 transplants. **(G)** Tumor growth curves showing tumor volume over time after tumor onset. Three biologically independent experiments were combined. WT, N=20; Hom, N=32; Het, N=32 transplants. **(H)** Quantification of total flux (luminescence) and representative bioluminescence images for lung metastasis at 8 weeks post-transplant. N=12 mice. Three biologically independent experiments were combined.

To functionally evaluate whether *KMT2C*/*MLL3* is haplo-insufficient in suppressing breast tumorigenesis, we generated a MLL3 heterozygous deletion breast tumor model (MLL3 Het) using the genetically engineered mouse mammary stem cell (MaSC) organoid-based breast tumor model approach we developed^[15]^. It is reported that the MLL3 SET domain is required for protein stability^[16]^, and we previously demonstrated that SET domain deletion caused severe MLL3 protein degradation in MaSCs^[9]^. Therefore, we utilized MaSCs from MLL3 SET domain floxed mice^[16]^ to generate MLL3 Het and Hom loss breast tumor model (**Fig. 1C**). Since *PIK3CA* activating mutation and *TP53* loss-of-function mutation are the most frequently co-occurring mutations in the *MLL3*-mutant human breast cancer^[9]^, we also introduced *Pik3ca* constitutive active mutation and *Trp53* gene deletion to recapitulate human patient gene mutation status as shown in our previous study^[9]^. The SET domain floxed mice were crossed with *Pik3ca^H1047R^* mice to generate MLL3^+/+^, MLL3^flox/+^, and MLL3^flox/flox^ mice with LoxP-Stop-LoxP-*Pik3ca^H1047R^*. MaSCs were then isolated and transduced with lentiCRISPRv2 virus to introduce Trp53 gene deletion by CRISPR/Cas9 and then Adeno-Cre virus to delete the floxed alleles (**Fig. 1C**). The *Trp53* deletion and PIK3CA activation were validated by western blot (**Fig. 1D**). The MLL3 protein was significantly reduced by MLL3 SET domain Hom deletion (KO), and the Het deletion model exhibited the intermediate level of MLL3 protein, mimicking het loss (**Fig. 1E**).

The resulting MLL3-WT, -KO, and -Het MaSCs (also carrying *Trp53*-KO and *Pik3ca^H1047R^*) were then transplanted into mammary fat pads of WT FVB recipient mice to monitor their tumorigenic activity. As observed in our previous studies^[9]^, MLL3-KO MaSCs initiated tumors significantly earlier and grew faster than MLL3-WT MaSCs (**Fig. 1F-G**). Upon injected through tail vein, the MLL3-KO MaSCs formed greater lung metastasis **(Fig. 1H**). Interestingly, the MLL3-Het model also showed a faster tumor onset than WT, with an onset speed between MLL3-WT and MLL3-KO (**Fig. 1F**), suggesting the haplo-insufficiency of MLL3 in suppressing breast tumor initiation. Beyond tumor onset, MLL3-Het tumors did not grow faster or form more metastasis than MLL3-WT (**Fig. 1G, H**), suggesting greater loss of MLL3 function is needed for driving these later events.

### Histone methyltransferase activity of MLL3 is not required for breast tumor suppression

While one prominent function of MLL3 is as a histone methyltransferase catalyzing histone H3 lysine 4 mono- and di-methylation, MLL3 also functions as a nuclear adaptor protein by recruiting other epigenetic regulators such as BAP1 and TET3^[12, 13]^. Since previous studies demonstrated tumor suppressor function of MLL3 using gene deletion approaches resulting in loss of the entire protein^[6, 7, 9, 12, 17]^, which MLL3 function is important for tumor suppression remains unclear. To dissect domain-specific functions of MLL3 for breast tumor suppression, we established the MLL3 histone methyltransferase catalytic dead mutant breast tumor model. The Y4792A point mutation within the SET domain (**Fig. 2A**), which is responsible for MLL3’s catalytic activity, has been reported to impair MLL3’s histone catalytic activity^[14]^. To establish a catalytic dead mutant model, MLL3^Y4792A^ transgenic mice^[18]^ were crossed with Pik3ca^H1047R^ mice as in the MLL3 Het loss model preparation. The MaSCs from MLL3^Y4792A^ / Pik3ca^H1047R^ mice were subjected to a series of virus transductions to delete the Trp53 gene and activate Pik3ca^H1047R^ transgene (**Fig. 2A, 2B**). MLL3^Y4792A^ mutant MaSCs showed a significant reduction of H3K4me1 (**Fig. 2C**), although other H3K4 methyltransferase may compensate for the global H3K4me1 level.

**Figure 2.**
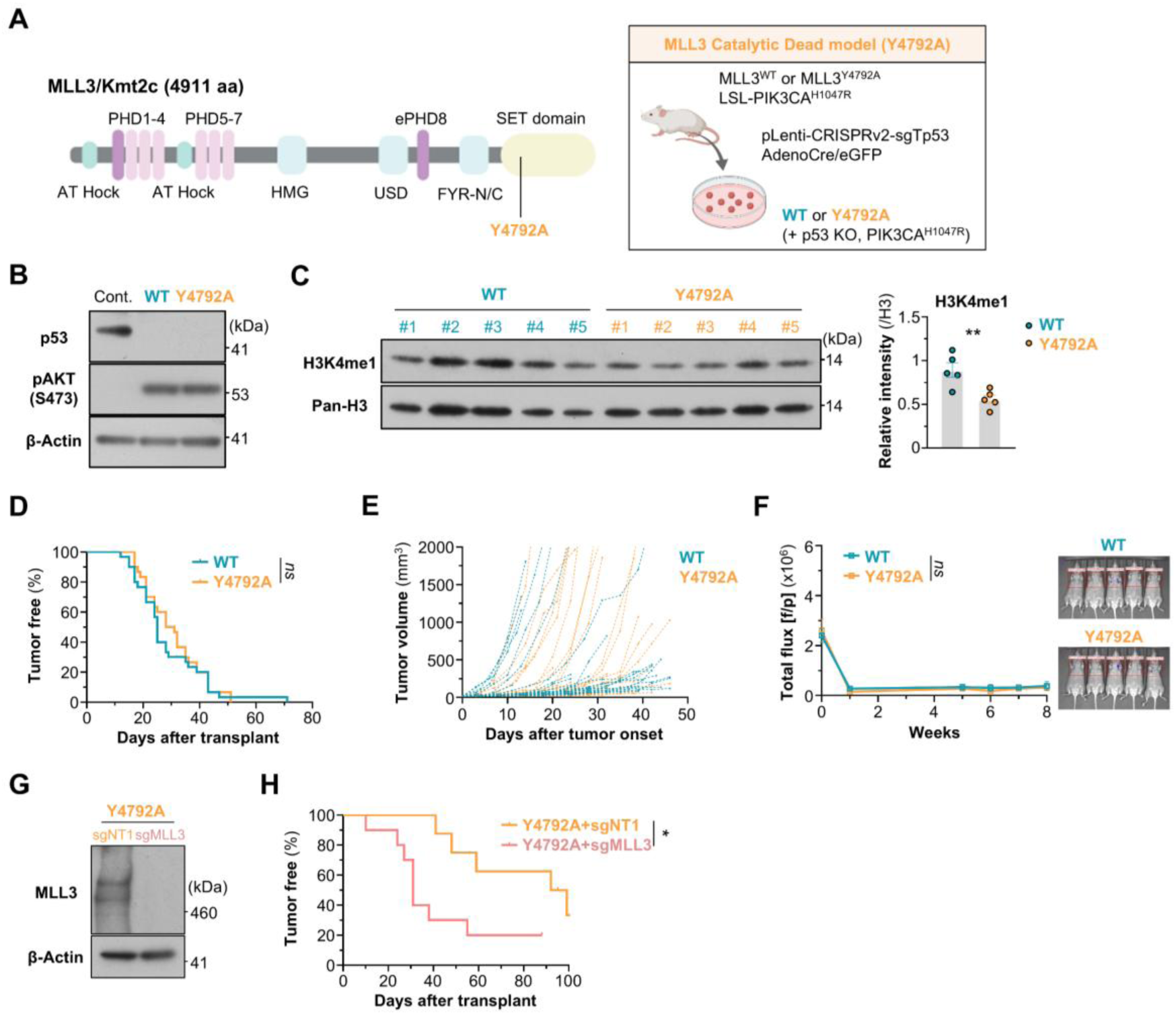
The histone methyltransferase catalytic activity of MLL3 is dispensable for breast tumor suppression. **(A)** Schematic overview of the generation of the MLL3 catalytic dead (Y4792A) mutant model. **(B)** WB analysis of Trp53 and pAKT (S473) in MLL3 WT and Y4792A mutant MaSCs. β-Actin was used as an internal control. **(C)** WB analysis of global H3K4me1 levels in MLL3 WT and Y4792A MaSCs. The right graph shows relative intensity of H3K4me1 normalized to that of pan-H3. Five biologically independent MaSC pairs were subjected to this analysis. **(D)** Tumor onset curve of MLL3 WT or Y4792A mutant MaSC transplanted mice. Three biologically independent experiments were combined. N=30 transplants/group. **(E)** Tumor growth curves showing tumor volume over time after tumor onset. Three biologically independent experiments were combined. WT, N=30; Y4792A, N=28 transplants. **(F)** Quantification of total flux (luminescence) and representative bioluminescence images for lung metastasis at 8 weeks post-transplant. N=10 mice/group. Two biologically independent experiments were combined. **(G)** WB analysis of MLL3 in sgNT or sgMll3 gRNA-expressing Y4792A MaSCs. β-Actin was used as an internal control. **(H)** Tumor onset curve with sgNT or sgMll3 gRNA-expressing MLL3 Y4792A breast tumor models. Representative result from two biologically independent experiments was shown. sgNT=8; sgMll3=10 transplants.

To evaluate their tumorigenesis activity, MLL3^WT^ and MLL3^Y4792A^ MaSCs were transplanted to the mammary fat pad of WT recipient mice. In contrast to MLL3 Hom deletion, MLL3^Y4792A^ exhibited a similar tumor onset and growth rate as MLL3^WT^ and failed to form macro-metastasis into the lung (**Fig. 2D-F**). To further test the tumor suppressing function of MLL3^Y4792A^, we further deleted the MLL3 gene in MLL3^Y4792A^ MaSCs by CRISPR, which led to significant acceleration of tumor onset (**Fig. 2G, H**). Taken together, the results demonstrated that the catalytic activity of MLL3 is not required for the tumor suppressor function of MLL3.

### The PHD2 domain-mediated adaptor function of MLL3 is indispensable for breast tumor suppression

The above results suggest that MLL3’s methyltransferase-independent function is important for breast tumor suppression. The PHD2 domain of MLL3 is responsible for binding with the BAP1 complex, which is an epigenetic regulator and known tumor suppressor, and is a hotspot mutation region in human cancer^[12, 19]^. Specifically, the G367V mutation in PHD2 (**Fig. 3A**) can inhibit the binding of MLL3 to the BAP1 complex^[12]^. Therefore, to test the MLL3 adaptor function in tumorigenesis, we generated MLL3^G367V^ MaSCs by utilizing a PE3 Prime Editing approach (**Fig. 3A**)^[20]^. Pik3ca^H1047R^ MaSCs were subjected to Prime Editing-based gene editing, followed by viral transductions to delete Trp53 gene and activate the Pik3ca^H1047R^ transgene. Two independent MaSC lines with 79 and 63% G367V conversion efficiency were established, as confirmed by amplicon high-throughput sequencing (**Fig. 3B**). Of note, structure analysis with AlphaFold2 showed no significant change in the overall structure around the PHD region by the G367V mutation (**Fig. S1A**). Consistent with the structural prediction, MLL3 protein level was comparable between MLL3^WT^ and MLL3^G367V^ mutant MaSCs (**Fig. 3C**), suggesting G367V point mutation does not affect MLL3 protein stability. The deletion of Trp53 and activation of Pik3ca transgene were validated by WB (**Fig. 3D**).

**Figure 3.**
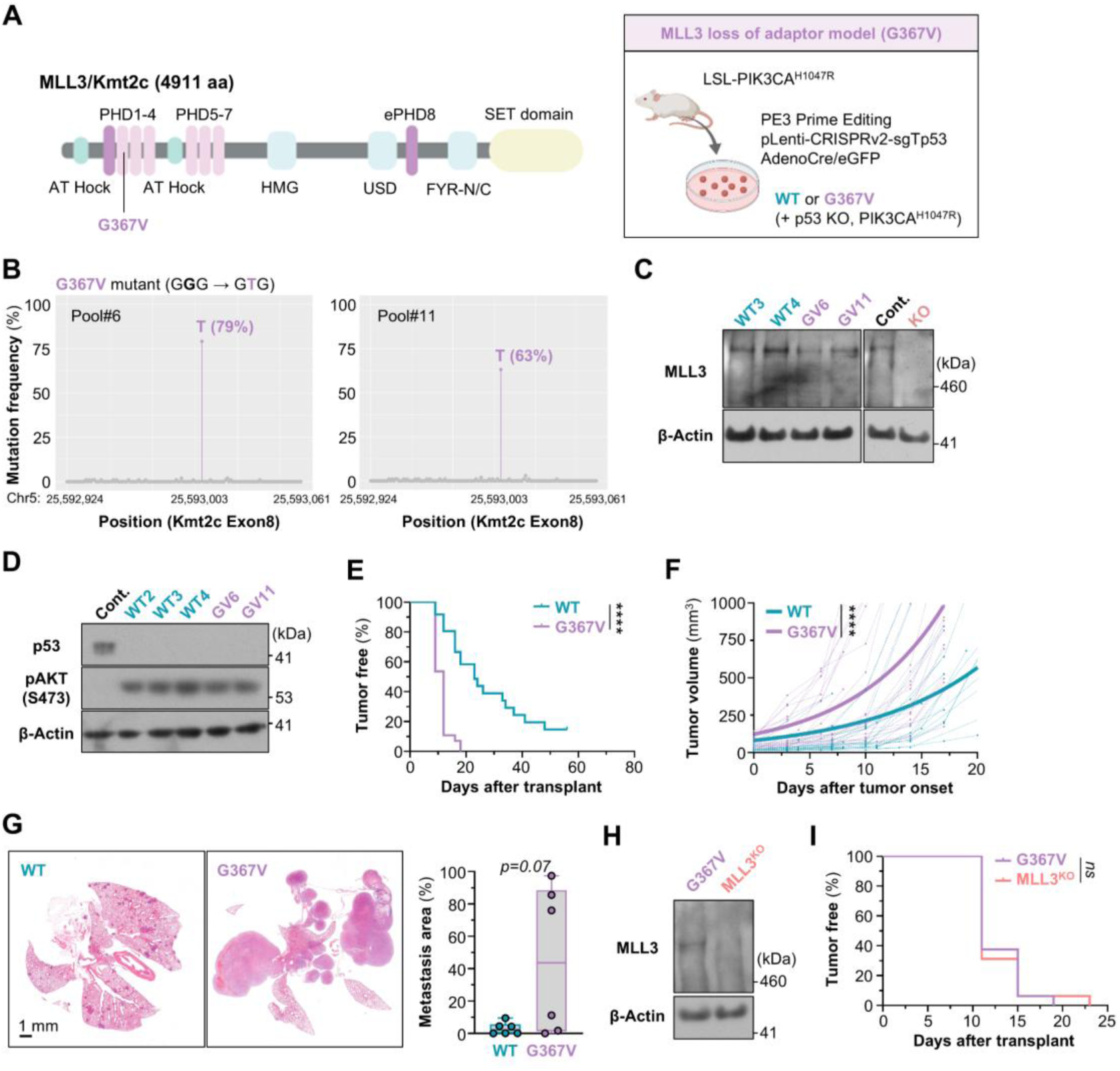
Loss of the PHD2 domain adaptor function of MLL3 phenocopies complete MLL3 loss. **(A)** Schematic overview of the generation of the MLL3 adaptor function loss (G367V) mutant model using PE3 Prime Editing. **(B)** Amplicon sequencing results determining the G-to-T mutation frequency in G367V MaSC lines. **(C)** WB analysis of MLL3 protein in MLL3 WT and G367V mutant MaSCs. β-Actin was used as an internal control. **(D)** WB analysis of Trp53 and pAKT (S473) in MLL3 WT and G367V mutant MaSCs. β-Actin was used as an internal control. **(E)** Tumor onset curve of MLL3 WT or G367V mutant breast tumor models. Two biologically independent experiments were combined. WT, N=30; G367V, N=28 transplants. **(F)** Tumor growth curves showing tumor volume over time after tumor onset. Two biologically independent experiments were combined. WT, N=29; G367V, N=28 tumors. **(G)** Representative H&E staining of lung sections showing metastases 8 weeks after tail vein injection. The right graph shows the percentage of lung area occupied by metastases. Two biologically independent experiments were combined. N=6 mice/group. **(H)** WB analysis of MLL3 levels in parental or sgMll3 gRNA-expressing G367V MaSCs. **(I)** Tumor onset curve of parental or sgMll3 gRNA-expressing G367V mutant breast tumor models. N=16 transplants/group.

In contrast to the catalytic dead mutant (MLL3^Y4792A^), the MLL3^G367V^ mutant MaSCs exhibited significantly faster tumor onset and growth and greater metastasis ability compared to MLL3^WT^ cells after transplantation (**Fig. 3E-G**). Moreover, MLL3 deletion by CRISPR in MLL3^G367V^ mutant MaSCs did not further accelerate tumor initiation ability (**Fig. 3H, I**), indicating that MLL3^G367V^ mutant phenocopies MLL3 Hom deletion. Of note, all of the tumor cells isolated from seven independent MLL3^G367V^ tumors showed complete G367V conversion (**Fig. S1B**), although the pre-implantation MLL3^G367V^ MaSCs (line#6) had only 79% conversion rate, suggesting a selection of homozygous MLL3^G367V^ mutation in breast tumors *in vivo*. Additionally, the expression of MLL3 protein was comparable between tumor-derived organoids from MLL3^WT^ and MLL3^G367V^ mutant tumors, further confirming that the G367V point mutation does not affect MLL3 protein stability (**Fig. S1C**).

We previously showed that MLL3 loss activates HIF1α signaling by stabilizing HIF1α protein stability, and the MLL3^KO^ tumor exhibited the unique immune suppressive tumor immune microenvironment (TIME) with an enrichment of highly differentiated regulatory T (T_reg_) cells^[9]^. Interestingly, the protein stability assay under pseudo-hypoxia conditions showed much greater stability of HIF1α in MLL3^G367V^ mutant MaSCs than WT (**Fig. S1D**), resulting in activation of the hypoxia gene signature (**Fig. S1E**). In addition, the spectral flow cytometry analyses with early-stage small MLL3^WT^ or MLL3^G367V^ tumors revealed the enrichment of highly differentiated T_reg_ cells, defined by high expression of effector T_reg_ cell markers, in MLL3^G367V^ mutant tumors (**Fig. S1F**).

Taken together, MLL3^G367V^ mutant tumors phenocopied MLL3^KO^ tumors, including faster tumor onset, tumor growth, greater HIF1α stability and enrichment of highly differentiated T_reg_ cells within the TIME.

### Loss of MLL3 impairs UTX chromatin localization and promotes breast tumor progression

Previous studies demonstrated that the binding of the MLL3 protein to the BAP1 complex could regulate chromatin localization of the MLL3 complex, including the UTX histone demethylase (**Fig. 4A**)^[12]^. We therefore hypothesized that the aggressive tumor phenotype of *MLL3*-mutant breast cancer may be mediated, at least in part, through altered BAP1 and UTX function. To test this possibility, we genetically deleted BAP1 or UTX in MLL3 WT MaSCs using CRISPR. Indeed, the deletion of either BAP1 or UTX also accelerated breast tumor initiation (**Fig. 4B-D, S2A, S2B**) with UTX depletion showing the stronger effect. Interestingly, level of UTX protein in nuclear chromatin fraction was markedly reduced by MLL3 loss in MaSCs (**Fig. 4E**) even though total protein levels of UTX and BAP1 were not altered by either MLL3 loss or G367V point mutation in MaSCs (**Fig. S2C**). Consistent with these findings in the mouse model, using published ChIP-seq data^[3]^, we found that MLL3 depletion in human breast cancer cell lines also showed a reduction in genome-wide occupancy of UTX (**Fig. 4F, S2D**). These results suggest that impaired chromatin localization of UTX may mediate the effect of MLL3 loss in promoting tumor onset and growth. To further test this mechanism, we further deleted UTX in MLL3 KO MaSCs using CRISPR. In contrast to the strong effect observed in MLL3 WT MaSCs, UTX depletion in MLL3 KO MaSCs did not further accelerate their tumor onset (**Fig. 4G, S2E**), suggesting that UTX mainly functions through the MLL3 complex in breast tumor suppression.

**Figure 4.**
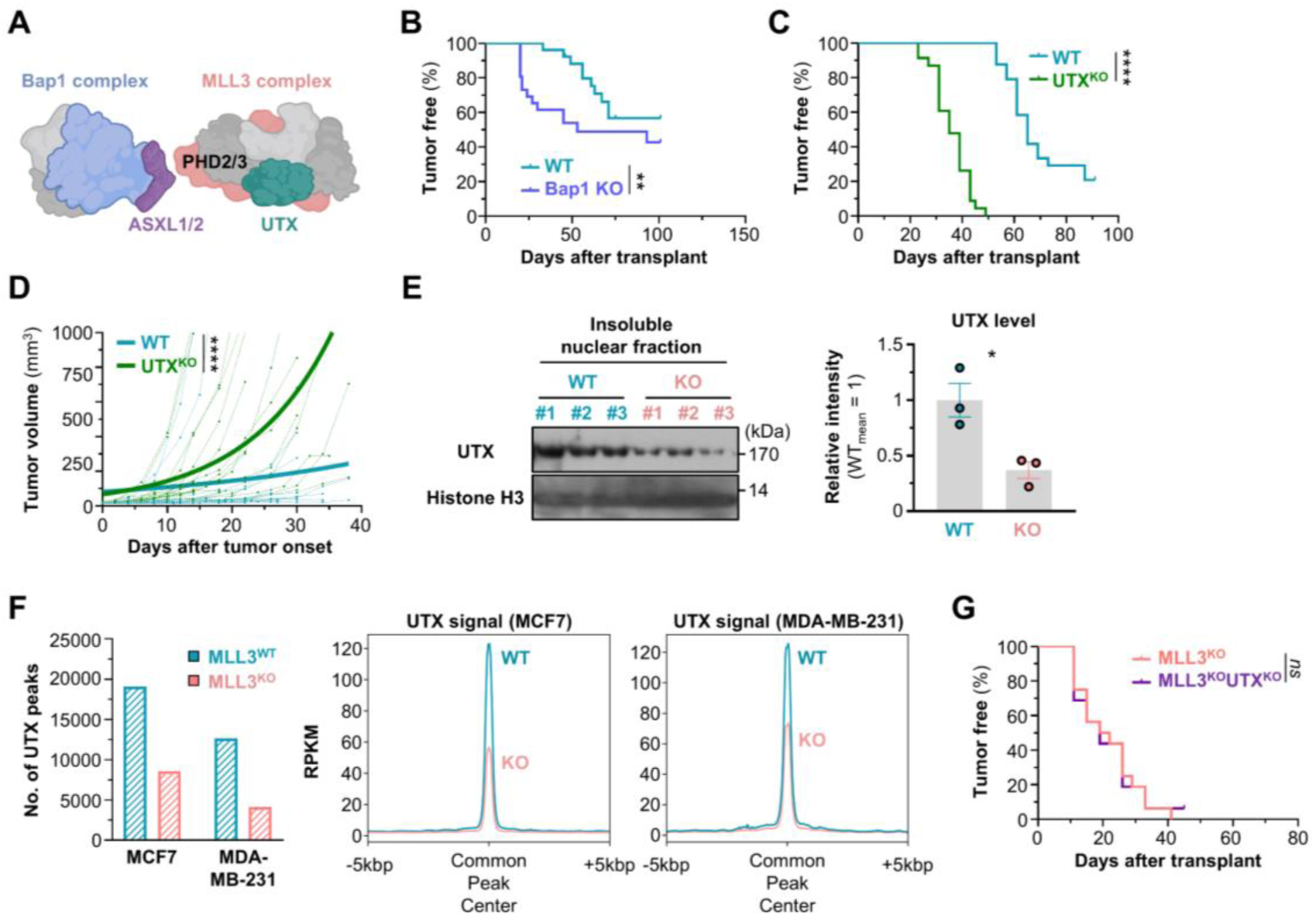
Loss of MLL3 impairs UTX chromatin localization and promotes breast tumor progression. **(A)** Schematic diagram of the binding of the MLL3 complex, including UTX, with the BAP1 complex. **(B)** Tumor onset curve of sgNT or sgBap1 gRNA-expressing MLL3 WT breast tumor models. Two biologically independent experiments were combined. N=26 transplants/group. **(C)** Tumor onset curve of sgNT or sgUTX gRNA-expressing MLL3 WT breast tumor models. Two biologically independent experiments were combined. sgNT, N=24; sgUTX, N=23 transplants. **(D)** Tumor growth curves showing tumor volume over time after tumor onset. Two biologically independent experiments were combined. sgNT, N=19; sgUTX, N=23 transplants. **(E)** WB analysis of UTX levels in the chromatin fraction of MLL3 WT and KO MaSCs. Pan-H3 histone was used as an internal control. The right graph shows relative intensity of UTX normalized to the average intensity of UTX in WT group. N=3 MaSCs/group. **(F)** UTX ChIP-seq data analysis in MLL3 WT and KO human breast cancer cell lines. The left graph shows number of UTX peaks. The right plots show the UTX signal around the common peak centers between MLL3 WT and KO cell lines. RPKM, Reads Per Kilobase of transcript per Million mapped reads. **(G)** Tumor onset curve with sgNT or sgUTX gRNA-expressing MLL3 KO breast tumor models. N=16 transplants/group.

### Epigenetic alterations at promoter regions, not enhancer regions, caused by MLL3 loss are correlated to gene expression changes in both MLL3 KO and G367V MaSCs

Considering the functions of MLL3 and UTX as epigenetic regulators, MLL3 loss could confer an aggressive breast tumor phenotype via changes in epigenetic landscapes coupled with gene expression changes. To address this point, the global transcriptome, histone modification, and chromatin accessibility in MLL3 KO MaSCs were compared to MLL3-WT cells by RNA-seq, CUT&TAG, and ATAC-seq, respectively. Transcriptome in MLL3^G367V^ mutant MaSCs was also analyzed, shown to share significant numbers of differentially expressed genes in MLL3 KO cells, including “acute-phase response” signature defined by serum amyloid A genes (e.g. *Saa1*) (**Fig. S3A-C**). For CUT&TAG analysis, H3K4me1 (catalyzed by MLL3), H3K27me3 (removed by UTX), and H3K27Ac (catalyzed by CBP/p300, MLL3 binding protein) marks were examined since MLL3 and its binding proteins regulate these histone modifications.

As expected, MLL3 loss reduced in the number of H3K4me1 and H3K27Ac peaks even though there were no major changes in the distribution pattern (**Fig. 5A**). In contrast, the number of H3K27me3 peaks was comparable between MLL3 WT and KO MaSCs. Differential peaks were also identified by the Diffbind pipeline, and consistent with the number changes, the majority of H3K27Ac differential peaks were downregulated peaks in MLL3 KO (**Fig. 5B**). Next, to evaluate putative enhancer activities regulated by MLL3, we defined “WT active enhancer” regions as genomic regions where H3K4me1 unique peaks overlapped with upregulated H3K27Ac differential peaks in MLL3 WT cells, or “WT poised enhancer” regions as genomic regions where H3K4me1 unique peaks overlapped with upregulated H3K27me3 differential peaks in MLL3 WT cells. Interestingly, the expression of the nearest genes from the putative WT unique active/poised enhancer regions were not markedly changed compared to all genes as background references (**Fig. 5C**). On the other hand, the genes with differential peaks of H3K27me3 (repressive) or H3K27Ac (active) around their promoter regions exhibited bigger changes in their gene expression, and the directions of the changes were consistent with the function of histone modifications (**Fig. 5C**). Of note, the expression of these genes with the unique epigenetic changes in their promotor regions were also affected by the MLL3^G367V^ mutation as in MLL3-KO cells (**Fig. 5D**). Moreover, chromatin accessibilities, examined by ATAC-seq, around promoter regions of the genes identified by CUT&TAG analysis (**Fig. 5C**) were correlated to their histone modification status with bigger effect on genes with repressive histone marks in MLL3 KO (**Fig. 5E**). Number of differentially more open peaks (9133 peaks) in MLL3 KO is less than that of close peaks (12074 peaks) (**Fig. 5F**), suggesting an increase in closed chromatin regions in MLL3 KO cells. Motif analysis with differential peaks identified binding motifs for TEAD and TP53/63/73 in MLL3 KO open and close regions, respectively, even though AP-1 binding motifs were enriched in both regions (**Fig. 5F**). In addition to the differential peak analysis, reproducible peak analysis also indicated MLL3 loss led to reduced chromatin accessibility, especially in intergenic regions (**Fig. 5G**). Motif analysis with WT- and KO-unique peaks further suggested that 1) AP-1 motif was highly enriched in both regions, and 2) TEAD and RUNX motifs were enriched in KO- and WT-unique regions, respectively, while KLF family motifs (most similar to KLF4 motif) were enriched in KO-unique regions (**Fig. 5H**).

**Figure 5.**
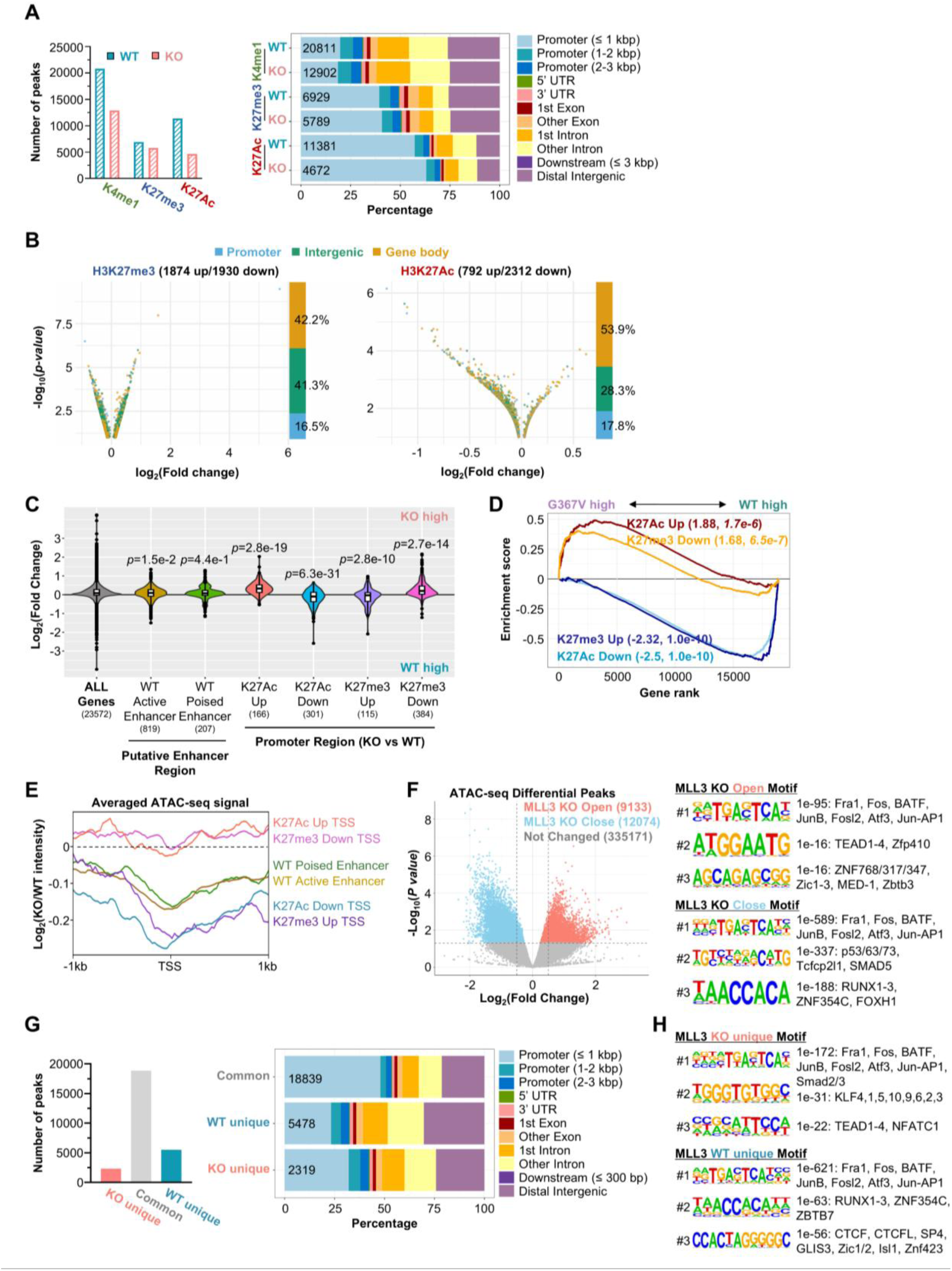
MLL3 loss induces aberrant gene expression primarily through epigenetic alterations at promoter regions. **(A)** Number of reproducible peaks (left) and genomic distribution (right) of H3K4me1, H3K27me3, and H3K27Ac in MLL3 WT and KO MaSCs determined by CUT&TAG. N=6 MaSCs/group. **(B)** Volcano plots highlighting differential peaks and their genomic distribution (Promoter, Intergenic, Gene body) for H3K27me3 and H3K27Ac between MLL3 WT and KO MaSCs. **(C)** Violin plots showing gene expression changes associated with putative enhancer and promoter regions with specific epigenetic alterations. The expression of the genes nearest to the annotated genomic regions was plotted. **(D)** Gene Set Enrichment Analysis (GSEA) plot showing that the genes with altered epigenetic landscapes in their promoter regions in MLL3 KO MaSCs, as defined in C, are regulated with the same directionality in MLL3 G367V mutant MaSCs. WT, N=3; G367V, N=2 MaSCs. **(E)** Chromatin accessibility around promoter regions of genes of various histone modification changes identified by CUT&TAG. The log2-scale (KO/WT) averaged bigwig files of ATAC-seq (N=6 MaSCs/group) were visualized around promoter regions of genes identified by CUT&TAG analysis in C. **(F)** Differential ATAC-seq peaks (p value < 0.05) called by DESeq2 were visualized as a volcano plots (left). The motif analysis (right) was performed by Homer, and top3 de novo motifs and predicted binding transcription factors (TFs) are listed based on p value. **(G)** Number of reproducible peaks (left) and peak annotation by ChIPseeker (right) in MLL3 WT and KO MaSCs. N=6 MaSCs/group. **(H)** Motif analysis with WT- or KO-unique peak regions. Top3 de novo motifs and predicted binding TFs are listed based on p value.

Taken together, the integrated analysis with RNA-seq, CUT&TAG, and ATAC-seq indicated that histone modification alterations in the promoter regions caused by MLL3 loss is more correlated with the gene expression and chromatin accessibility changes than histone modification changes in the putative enhancer regions. The genes with altered histone modifications in their promoter regions in MLL3 KO MaSCs were also similarly affected by MLL3^G367V^ mutation, supporting the importance of nuclear adaptor function of MLL3 in controlling gene expression.

## Discussion

In this study, we identify that the breast tumor suppressor function of MLL3/KMT2C is mainly mediated by its catalytic-independent adaptor function of MLL3/KMT2C rather than its histone methyltransferase function. Using genetically defined MLL3 loss breast tumor models, we show that MLL3 suppresses breast tumor initiation in a dosage-sensitive manner. Furthermore, the mutation of MLL3 PHD2 domain that disrupts its binding to the BAP1 complex phenocopies MLL3 homozygous loss, whereas the mutation inactivating MLL3 methyltransferase has no effect on tumor initiation or growth.

Although MLL3 is frequently mutated in various types of human cancers^[2, 21]^, how heterozygous loss of MLL3 contributes to tumor progression has remained incompletely understood. In neurodevelopmental disorders, KMT2C haploinsufficiency has been established through large-scale human patient cohort and mouse genetic studies^[22–24]^, indicating that MLL3 function is dosage sensitive in specific biological contexts. In the cancer context, however, the studies about MLL3 dosage effects have been more limited. In acute myeloid leukemia (AML), partial reduction of MLL3 expression using RNAi- and CRISPR-based approaches was shown to accelerate leukemogenesis in a haplo-insufficient manner, supporting a haploinsufficiency of MLL3 for tumor suppression in AML^[6]^. However, whether MLL3 acts as a dosage-sensitive tumor suppressor in solid tumors has remained poorly defined. Our MLL3 SET domain floxed breast tumor model demonstrates that partial reduction of MLL3 protein is sufficient to accelerate tumor initiation, although it does not promote later tumor growth or metastasis in this model. Importantly, the majority of MLL3 mutations in breast cancer patients are heterozygous deletion^[3]^. These findings suggest that monoallelic KMT2C alterations may be positively selected during early breast tumor development by lowering the threshold for tumor initiation, even if additional genetic or epigenetic events, such as transcription downregulation of MLL3, are required for more advanced tumor progression.

An unexpected finding of our study is that the catalytic activity of MLL3 is dispensable for its tumor-suppressive function in our breast tumor model context. The Y4792A mutation in the SET domain of MLL3, which impairs the histone methyltransferase activity, did not accelerate tumor onset, growth, or metastasis. Moreover, further deletion of the Y4792A mutant MLL3 allele in the MLL3^Y4792A^ MaSC accelerated tumor formation, indicating that the presence of the MLL3 protein, rather than its catalytic activity per se, is critical for suppressing breast tumor progression. Interestingly, previous studies with embryonic stem cells (ESCs) have reported that catalytic-dead MLL3/MLL4 mutants cause broad changes in H3K4me1 while producing relatively limited effects on gene expression compared to complete loss of MLL3 proteins^[14]^. In contrast, MLL3/MLL4 methyltransferase activities were later shown to control early embryonic development and ESC differentiation in a lineage-selective manner, while MLL3/4-catalyzed H3K4me1 was largely dispensable for enhancer activation during ESC differentiation^[18]^. Given the context-dependent manner of MLL3 enzymatic and adaptor functions, our findings revealed the key molecular activity of MLL3 in the context of tumor suppression. Our integrated transcriptomic and epigenomic analyses further suggest that MLL3-mediated transcriptional changes in breast cancer are more closely associated with promoter-proximal chromatin alterations rather than with canonical enhancer regulation. Although MLL3 loss reduced H3K4me1 and H3K27Ac peaks, genes linked to putative MLL3-mediated enhancer regions showed relatively modest expression changes. In contrast, genes associated with promoter-proximal changes in H3K27Ac or H3K27me3 displayed stronger and directionally consistent transcriptional alterations, and these gene sets were similarly regulated by the G367V mutation. These findings suggest that MLL3 loss may reshape tumor-promoting transcriptional programs through a primarily promoter-centered chromatin modification mechanism. Since putative enhancer-mediated genes in our analysis were inferred primarily by genomic proximity and distal enhancers can regulate genes located far from their linear genomic position, our data do not fully rule out a contribution of enhancer regulation to tumor phenotype of *MLL3*-mutant breast cancers.

In contrast to the catalytic-dead Y4792A mutation, the G367V mutation in the PHD2 domain phenocopied MLL3 loss. Importantly, the G367V mutation did not substantially reduce MLL3 protein abundance and deletion of the mutant MLL3 in G367V-mutant cells did not further accelerate tumor initiation, suggesting that this phenotype is not simply caused by protein destabilization. Given (1) the BAP1 gene deletion has been reported to impair chromatin localization of MLL3 and UTX proteins^[12]^, (2) both Bap1 and Utx loss accelerated breast tumor progression, (3) MLL3 loss markedly impaired UTX chromatin localization, and (4) UTX depletion promoted breast tumor progression in MLL3-proficient, not MLL3-deficient, setting, UTX could be the candidate downstream regulator in *MLL3*-mutant breast cancers. UTX is an H3K27me3 demethylase and known as a tumor suppressive chromatin regulator in several cancer contexts^[25–27]^. In breast cancer, prior studies have shown that UTX loss can induce cancer stem cell-like property via promoting epithelial-mesenchymal-transition (EMT), resulting in aggressive tumor progression^[28, 29]^. Of note, our previous work showed that MLL3 loss induces EMT-associated cellular plasticity and stem-like phenotypes in breast cancer cells^[3]^. This phenotypic overlapping between MLL3 deficiency and UTX loss further indicates MLL3 suppresses breast tumorigenesis, at least in part, by maintaining UTX-dependent chromatin regulation. Future studies integrating UTX occupancy, H3K27me3 dynamics, and transcriptional changes induced by MLL3 loss and G367V mutation will help define the epigenetic landscape controlled by the MLL3–BAP1–UTX axis and reveal epigenetic vulnerabilities created by disruption of this chromatin regulatory module in *MLL3*-mutant breast cancer.

A major strength of this study is the use of endogenous full-length MLL3 point-mutant models. Previous biochemical studies have mainly relied on short MLL3 protein fragments, or partial complexes ^[12, 13]^. Since MLL3 is an exceptionally large and structurally complex chromatin regulator, studying domain-specific functions using short protein fragments may not fully recapitulate the interactions of various domains in the full-length protein. By using prime editing to introduce the G367V mutation into endogenous MLL3, our study enabled functional interrogation of a specific regulatory interface while preserving the native protein architecture and expression context. This approach is particularly important for distinguishing true domain-specific functions from artifacts caused by protein truncation, instability, or non-physiological overexpression. Nevertheless, our study has several limitations. Although the G367V mutation was previously reported to impair the interaction between MLL3 and the BAP1 complex, we have not directly demonstrated that G367V disrupts BAP1 binding in the context of endogenous full-length MLL3. It also remains possible that this G367V mutation affects interactions with other proteins rather than BAP1 only. Unbiased proteomics approach to identify binding proteins of full-length MLL3 G367V mutant protein could provide further understanding of crucial chromatin regulator interactions mediated by MLL3.

In summary, our findings revealed MLL3 suppresses breast tumor initiation through a dosage-sensitive, catalytic-independent adaptor function. Rather than acting through its canonical H3K4 methyltransferase activity, MLL3 appears to coordinate BAP1/UTX-associated chromatin regulation to maintain tumor suppressive transcriptional states. These results provide a mechanistic framework for understanding how non-catalytic disruption of MLL3 can drive breast tumor progression and establish full-length MLL3 mutant models as powerful tools for defining the chromatin dependencies of *MLL3*-mutant breast cancers.

## Acknowledgements

We thank the Einstein FACS, Histopathology, Analytical Imaging, Computational Analysis and Stem Cell Cores. This work was supported by NIH grant R01CA212424 to WG. KN and MM received Paul S. Frenette scholar fellowship from Einstein Stem Cell Institute. Einstein core facilities were supported by the Montefiore Einstein Comprehensive Cancer Center (NCI cancer center support grant 2P30CA013330) and the NYSTEM shared facility (C029154).

## Author Contributions

KN designed, performed and interpreted most experiments, assembled figures, and contributed to writing/editing of the manuscript. JC and MM contributed to animal experiments. NC set up high-dimensional FACS panels. YM and MS advised CUT&TAG and ATAC-seq bioinformatics analysis. GL advised immune profiling experiment. GX and KG generated MLL3 mutant mouse models and advised on the work. WG designed and interpreted experiments, contributed to figure design and editing, and wrote the paper. WG supervised the study.

## Materials and Methods

### Mice

All mice were bred in the SPF animal facility at the Albert Einstein College of Medicine. Wild-type (WT) FVB/n (JAX, #001800) and Pik3ca^H1047R^ (JAX, #016977) mice were purchased from the Jackson Laboratory. MLL3 SET domain floxed^[16]^ and Y4792A point mutant^[18]^ mice were provided by Dr. Kai Ge (NIDDK), and crossed with Pik3ca^H1047R^ mice. All animal experiments were carried out in strict accordance with the protocols approved by the Institutional Animal Use and Care Committee of the Albert Einstein College of Medicine. All efforts were made to minimize suffering and provide humane treatment to the animals included in the study.

### MaSC lines

Unless otherwise indicated, all MaSC lines used in this study carry Tp53 deletion and Pik3ca^H1047R^ mutation. MLL3 SET domain homozygous (Hom)/heterozygous (Het) loss, MLL3 Y4792A, MLL3 G367V, BAP1 KO, and UTX KO MaSCs were newly established in this study as described below. FVB/n background MLL3 Knockout (KO) MaSC carrying Tp53 deletion and Pik3ca^H1047R^ mutation and counterpart MLL3 WT MaSC lines used in this study were established previously^[9]^.

### Isolating MaSCs and culturing MaSC organoids

Mammary glands from MLL3 WT or MLL3 mutant (SET-flox or Y4792A) Pik3ca^H1047R^ mice were minced and digested with 300 units/ml collagenase type 3 (Worthington Biochemical, LS004182), and 10 μg/mL Dnase I (Worthington Biochemical, LS002139) in the DMEM/F-12 medium at 37°C for 2 hours on an rotating mixer (100 units/mL hyaluronidase added only for tumor digestion). The digested samples were washed with PBS and were further digested with 0.05% trypsin-EDTA for 5 minutes and 1 unit/mL neutral protease (dispase) (Worthington Biochemical, LS02109) plus 100 μg/mL Dnase I (Worthington Biomedical) for 5 minutes. The digested cells were then filtered through 40 μm cell strainers to obtain single cells. To isolate tumor cells from established tumors (5-10 mm of diameters) for organoid culture, the tumors were minced as for mammary glands and digested with the same enzymes further supplemented with 100 units/mL hyaluronidase (Worthington Biochemical, LS002592), followed by the same isolation procedure.

Organoid culture was performed as previously reported ^[30, 31]^. Organoid medium was prepared with Advanced DMEM/F-12 (Life Technologies) supplemented with 5% heat-inactivated FBS (Sigma, F2442), 1x Glutamax supplement (Life technologies, #35050061), 10 ng/mL EGF (Sigma, E9644), 20 ng/mL FGF2 (EMD Millipore, GF003), 4 μg/mL heparin (Sigma, H3149), 5 μM Y-27632 (Cayman Chemical, #10005583) and 5% Matrigel (Corning, #354234). For passaging and expansion, organoids were washed with PBS, dissociated with 0.05% trypsin/EDTA and seeded at 3 × 10^5^ cells/well in 6-well ultra-low attachment plates. Cells were then passaged every 3-4 days. Dissociated organoid cells were frozen in bovine calf serum containing 10% DMSO and 5 μM Y-27632, and thawed to reestablish the culture.

### Establishment of genetically engineered MaSC

#### Viral transduction

Lentiviruses were produced with the pMD2.G and pCMVR8.74 packaging system, a gift from Didier Trono (Addgene, plasmids #12259 and #22036), in HEK293T cells. Viruses were concentrated with the Lenti-X Concentrator reagent (Clontech, #631232). For viral transduction, dissociated organoid cells were seeded in virus-containing organoid culture media with 5 μg/mL polybrene (EMD-Millipore TR-1003-G) overnight, and medium was refreshed the next day. Transduced cells were either FACS sorted by GFP or RFP or selected by puromycin (2 μg/mL) treatment.

#### CRISPR-based genome editing

Genome editing for MLL3 mutant (SET domain Het/Hom floxed or MLL3 Y4792A) / Tp53^null^ / Pik3ca^H1047R^ MaSCs (FVB/n background) were performed as following: MaSCs isolated from MLL3 mutant Pik3ca^H1047R^ mice were transduced with sgTP53 cloned into lentiCRISPRv2-puro (Addgene, #98290). After one passage, cells were grown for 3 days and selected with puromycin for 3 days. Next, cells were infected with Adeno-Cre-GFP and sorted for GFP positive cells. BAP1, UTX, and MLL3 KO lines were made via further transducing the sorted MaSCs with sgBap1, Utx, Mll3 or sgNT cloned into the modified pLKO5.sgRNA.EFS.tRFP (Addgene #57823, for Bap1), which accept two distinct guide RNAs, or lentiCRISPRv2-GFP (Addgene, #82416, for Utx and Mll3) and sorting for RFP or GFP. Targeting sequences for sgRNAs are as following: sgMll3 GCACACGATCTAGTACTCAG (Exon10), sgTp53 CCTCGAGCTCCCTCTGAGCC (Exon 2), sgNT GCGAGGTATTCGGCTCCGCG or GCTTTCACGGAGGTTCGACG, sgBap1 GGCGGTGGGCGCTCCCGGTT (Exon 1) and ATCCTTCATTCGGCTCAGCG (Exon 5), and sgUtx CTGGTATGCAGATAATGCTA (Exon 4).

Genome editing for MLL3 G367V / Tp53^null^ / Pik3ca^H1047R^ MaSCs (FVB/n background) were performed as following: MaSCs isolated from Pik3ca^H1047R^ mice were transiently transfected with three PE3 plasmids, including 1) pU6-pegRNA-GG-acceptor (Addgene, #132777) inserted with pegRNA, scaffold, and extension sequences, 2) pCMV-PE2-P2A-GFP (Addgene, #132776), and 3) pLKO5.sgRNA.EFS.tRFP inserted with PE3 sgRNA, using NEPA21 Electroporator (Bulldog Bio). Three days after transfection, the GFP/RFP double positive cells were sorted and expanded by further organoid culture. Point mutation was confirmed by Sanger and Amplicon sequencing with genomic PCR amplicon using the primers as followed; Forward AGCCCGGGAGATCTTTTAGA, Reverse GGCCAAGGGATGAAATACAA. Amplicon sequencing was performed by MGH CCIB DNA Core, and the Fastp-processed amplicon sequencing reads were aligned to the mouse GRCm39 primary assembly reference genome using bwa-mem2. SAM files were converted to BAM format, followed by BAM sorting and indexing using SAMtools. The point mutation rate was visualized and examined via IGV browser. Once the point mutation was validated, the mutant MaSCs were further subjected to Trp53 deletion and Pik3ca^H1047R^ transgene activation as described above for the other lines. The sequences for oligos used in Prime Editing were listed in **Table S1**. Predicted structural models of the WT and G367V MLL3 PHD2 region were generated using ColabFold^[32]^, an AlphaFold2-based protein structure prediction pipeline, and visualized in UCSF ChimeraX^[33]^.

### Tumor onset, growth, and metastasis assay

Organoids were cultured for 3-4 days, then dissociated and passed through 40 μm cell strainer to obtain single cells. Single cells (3 x 10^5^) suspended in PBS and 25% Matrigel were orthotopically injected into #2/3 or #4 mammary gland of FVB background female recipient mice aged 6-8 weeks. For tumor onset assay with UTX and MLL3 DKO and MLL3 KO in G367V backbone, 1 x 10^5^ cells per gland were transplanted. For tumor onset and growth experiments, four glands per mouse were injected. Tumor onset was monitored by weekly palpation, and Kaplan-Meier survival analysis was used to compare tumor onset. Tumor onset was scored when at least one dimension of the tumor measured ≥3mm. Tumor volume was measured by caliper and calculated as V = 0.532 × Length × Width^2^. For metastasis assay, 5 x 10^5^ single cells suspended in PBS were injected intravenously via the tail vein. To monitor the metastasis growth, the injected MaSCs were transduced with pCDH-EF1-Luc2-P2A-tdTomato (Addgene,#72486) for IVIS analysis. For the IVIS monitoring, the mice were intraperitoneally injected with D-luciferin (150 mg/kg body weight; GoldBio, LUCNA-100) 15 minutes before bioluminescence imaging. Formalin-fixed paraffin-embedded (FFPE) sections for the lung eight weeks after injection were subjected to Hematoxylin/Eosin staining to measure the metastasized area by ImageJ. Histological staining slides were scanned by 3DHistec Pannoramic 250 Flash II slide scanner.

### Western blot

Western blots were performed with antibodies against MLL3, phospho-Akt (Ser473), HIF1α, H3K4me1, Histone H3, Tp53, BAP1, UTX, and β-actin (see Table S1 for antibody sources and dilutions). Whole cell lysate proteins were isolated using RIPA-based isolation protocol described previously^[34]^. For detecting MLL3 protein, the proteins were isolated using NP-40 based mild lysis reagent optimized for MLL3/4 protein^[35]^. To reduce non-specific binding of anti-MLL3 antibody, pre-absorbed anti-MLL3 antibody was used as described previously^[9]^. Briefly, MLL3 KO MaSCs were seeded in a Nunc Lab-Tek II Chamber Slide (Thermo Scientific, #154453) without Matrigel and cultured under adherent conditions. Two days after seeding, when cells reached approximately 95% confluency, the cells were washed with PBS and fixed with 4% PFA. Fixed cells were permeabilized with 0.2% Triton X-100 in PBS and blocked with blocking buffer consisting of 2% BSA, 5% normal goat serum, and 0.1% Tween-20 in PBS. The cells were then incubated overnight with anti-MLL3 antibody (1:100) diluted in staining buffer consisting of 5% BSA, 5% normal goat serum, and 0.1% Tween-20 in PBS. The antibody-containing solution recovered after this incubation was further diluted (1:1000) to use as the pre-absorbed anti-MLL3 antibody for western blotting. For HIF1α stability test, MLL3 WT and G367V MaSCs were cultured in the presence of DMOG (1 mM) to accumulate HIF1α protein, following with the cycloheximide treatment (30 µg/mL) for 0 to 8 hours. For detecting UTX protein in chromatin fraction, protein fractionation from the cell pellets was performed using LysoPure Nuclear and Cytoplasmic Extractor Kit (FUJIFILM Wako Pure Chemical Corporation, 295-73901) by following manufacturer’s instructions.

### Regulatory T cell profiling using Spectral flow cytometry

Pre-onset MLL3 WT (day 20) and G367V (day 7) mutant tumors (1-3 mm diameter) were minced into 1 mm fragments, which were incubated in PBS with 10 UI/mL collagenase I, 400 UI/mL collagenase D and 30 µg/mL DNase I at 37°C for 1 hour under agitation. After, the fragments were homogenized, filtered through a 100 µm nylon mesh cell strainer and cells were centrifuged at 700 x g for 15 minutes at 8°C to obtain cell suspensions. Cell suspensions were incubated with CD16/32 blocking antibodies (BioLegend, # 101301, Clone:93) in FACS buffer consisting of 0.5% BSA, 2 mM EDTA, 0.1% sodium azide in PBS and stained with fluorescently tagged Abs purchased from eBioscience, BD Biosciences, R&D systems, or BioLegend (See **Tables S2**) diluted in FACS buffer. Brilliant stain buffer (BD) was also used for BD Horizon Brilliant fluorescent polymer dyes. Cells were stained for cell-surface marker expression, then fixed in eBioscience Fixation/Permeabilization buffer prior to intracellular Foxp3 transcription factor (TF) staining in eBioscience Permeabilization buffer for 1 hour. Data acquisition was done using a Cytek Aurora flow cytometer. All flow cytometry data were analyzed using FlowJo v10 software (BD Biosciences).

### Public database analyses

For analyses of overall survival and KMT2C mRNA expression of KMT2C WT, Het loss, and Hom loss patients, the dataset from the TCGA BRCA cohort (PanCancer Atlas, 1084 patients) was analyzed via cBioPortal.

### Epigenome and Transcriptome analysis

#### CUT&TAG analysis

To profile genome wide distribution of histone modifications, CUT&TAG analysis targeting H3K4me1, H3K27me3, and H3K27Ac was conducted by following EpiCypher’s CUT&Tag Protocol (v1.6). Briefly, 1 x10^5^ dissociated MLL3 WT or KO MaSCs were subjected to nuclei extraction, and the extracted nuclei were mixed with pre-activated binding beads. Then, the mixture of the nuclei and beads was incubated with primary antibodies at 4°C overnight. After washing with Digitonin buffer, CUTANA pAG-Tn5 enzyme was added, followed by the incubation at 37°C for 1 hour for chromatin tagmentation. The tagmented chromatin was subjected to library preparation by PCR using CUTANA High Fidelity PCR Master Mix and barcode-containing primers (see **Table S1**). The PCR products were purified with AMPure XP beads, and the purified DNAs were processed for next-generation sequencing using NovaSeq PE150 (Novogene).

For the CUT&TAG data analysis, first, the sequencing reads were processed using fastp program^[36]^ for quality control and trimming of low-quality reads. The cleaned reads were aligned to mm10 mouse genome using BWA-MEM2 algorithm^[37]^. After removing the reads that were duplicated or mapped to mitochondrial chromosomes using Picard^[38]^, the aligned reads were used to identify significant peaks with the MACS2 algorithm^[39]^ (H3K4me1, K27me3 – broad peak, H3K27Ac – narrow peak) under the default setting compared with IgG controls. The peak files were further processed using ChIP-R and Diffbind programs^[40, 41]^ to identify reproducible and differentially enriched peaks, respectively. Finally, the peak regions were annotated using ChIPseeker program^[42]^. The regions where peaks of different histone modifications were overlapped were identified using BEDtools.

#### Transcriptome analysis

RNA-seq libraries were prepared from total RNA isolated from MLL3 WT, KO, and G367V mutant MaSCs using poly(A) enrichment and a non-directional mRNA-seq library preparation protocol. Libraries were sequenced on an Illumina NovaSeq 6000 platform with paired-end 150 bp reads. Raw FASTQ files were quality-filtered and pre-processed using fastp, and transcript abundance was quantified using Salmon against the mouse mm10 reference transcriptome. Gene-level count and TPM matrices generated by RaNA-seq^[43]^ were used for quality control, sample comparison, and downstream analyses. Differential gene expression analysis between groups was performed using DESeq2, and genes with adjusted p values less than 0.05 were considered differentially expressed genes. For gene set enrichment analysis, raw count data were normalized using VoomNormalize module in GenePattern, and the normalized expression matrix was used as input for GSEA with Hallmark gene sets, following parameters: Ratio_of_Class, Phenotype, Weighted. Custom gene set enrichment analysis was performed using clusterProfiler. Genes were ranked by log2 fold change from the RNA-seq differential expression analysis. Custom gene sets, including genes associated with increased or decreased H3K27Ac or H3K27me3 signals at TSS-proximal regions, were converted to Entrez IDs using org.Mm.eg.db and used as TERM2GENE inputs for GSEA.

#### ATAC-seq analysis

ATAC-seq libraries were generated based on the Omni-ATAC protocol with minor modifications^[44, 45]^. Briefly, 5 x 10^4^ MLL3 WT and KO MaSC cells were subjected to nuclei prep, followed by transposase reaction at 37°C for 30 minutes on thermomixer. Transposase inserted DNA were purified with Qiagen MinElute Reaction Cleanup column, and the purified DNA was subjected to library preparation using NEBNext High-Fidelity PCR enzyme and barcode-containing primers (see **Table S1**). The PCR product was further purified using Axygen AxyPrep MAG PCR Clean-up Kit. The DNA concentration and fragment size distribution were confirmed by qubit and bioanalyzer, and the ATAC-seq libraries were processed for next-generation sequencing using NovaSeq PE150.

Raw ATAC-seq reads were quality-filtered and adapter-trimmed using fastp. Cleaned reads were aligned to the mouse GRCm38 reference genome using BWA-MEM2. Uniquely mapped reads were retained, mitochondrial reads and duplicated reads were removed, and the resulting BAM files were restricted to standard chromosomes for downstream analyses. Peaks were called for each sample using MACS2 with default setting for narrowpeaks. Peaks overlapping the mm10 blacklist regions were removed using BEDtools. FRiP scores, defined as the fraction of mapped reads overlapping blacklist-filtered MACS2 peaks, were calculated using SAMtools and BEDtools for QC. A master peak set was generated from blacklist-filtered ATAC-seq peaks, and read counts over the master peak set were quantified from each BAM file using BEDtools multicov. Differential accessibility between MLL3 WT and KO samples was analyzed using DESeq2 and the master peak matrix. For visualization, normalized bigWig files were generated from final BAM files using bamCoverage with RPGC normalization. Average WT and KO bigWig tracks were generated, and KO-versus-WT log2-ratio bigWig files were produced using bigwigCompare. ATAC-seq signals over selected genomic regions were summarized using computeMatrix and visualized using plotProfile. Reproducible peaks present in at least four of six replicates were identified using ChIP-R, and WT-unique, KO-unique, and common peak regions were further identified using BEDtools. ChIPseeker and HOMER were used for genomic region annotation and motif analysis, respectively.

#### Published UTX ChIP-seq data analysis

Published UTX ChIP-seq FASTQ files from MLL3 WT and KO MCF7 and MDA-MB-231 human breast cancer cell lines^[3]^ were downloaded and re-analyzed. Raw reads were quality-filtered and adapter-trimmed using fastp, followed by alignment to the human hg38 reference genome using Bowtie2. MACS2 was used for peak calling under default settings. Common and condition-specific UTX peaks between MLL3 WT and KO samples were identified using BEDtools intersect. Common peaks were defined as peaks overlapping between MLL3 WT and KO samples, whereas WT-only and KO-only peaks were defined as peaks detected in one condition but not overlapping peaks from the other condition. The resulting peak sets were annotated using ChIPseeker. For visualization, aligned BAM files were converted to FPKM-normalized bigWig files using bamCoverage. UTX ChIP-seq signal over common peak regions was summarized using computeMatrix and visualized using plotProfile.

## Statistical Analysis

P values for comparing the mean values of different groups were calculated using unpaired t-test except unless otherwise specified in figure legends. Log-rank test and Two-way-ANOVA was used in tumor onset and growth analysis, respectively. The peaks with FDR < 0.1 calculated by Diffbind program were identified as differential peaks in this study. All statistical analyses except for GSEA and omics analyses were done in GraphPad Prism 9 or 10 (*p< 0.05, **p< 0.01, ***p< 0.001, ****p< 0.0001, ns or not indicated: non-significant).

## Figure Legends

**Figure S1. Related to Figure 3.**
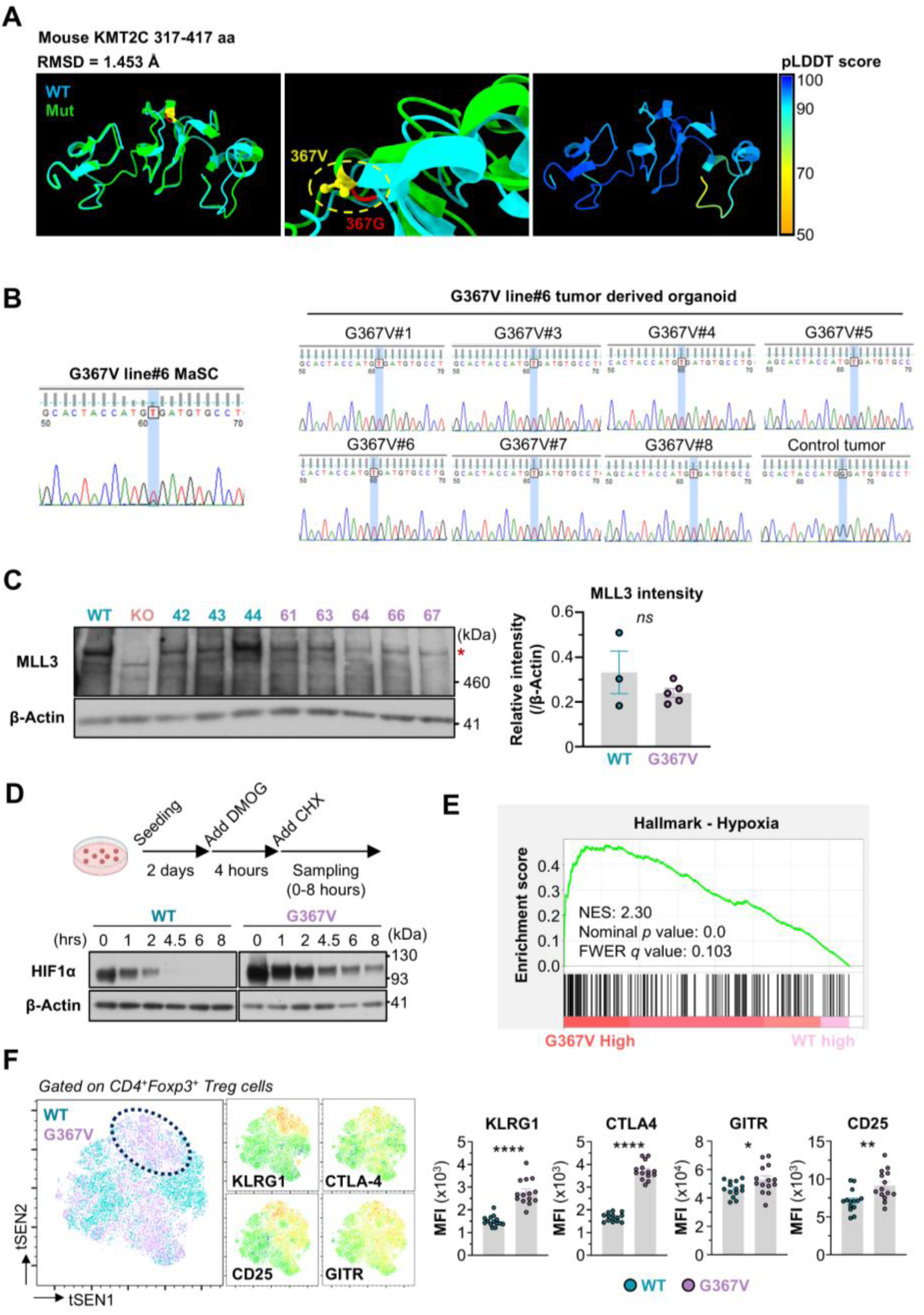
(A) Protein structural prediction of mouse MLL3 fragment around PHD2 region (aa 317-417) with or without G367V mutation by ColabFold, an AlphaFold2-based structure prediction pipeline. The structure with pLDDT scores of the G367V mutant peptide was illustrated on the right. RMSD, Root Mean Square Deviation. pLDDT, predicted Local Distance Difference Test. **(B)** Sanger sequencing showing the MLL3 G367V point mutation site in parental MLL3 G367V#6 MaSCs and independent tumor-derived organoids from MLL3 G367V#6 or WT tumors. **(C)** WB analysis of MLL3 protein levels in tumor-derived organoids from MLL3 WT and G367V tumors. β-Actin was used as an internal control. **(D)** HIF1α protein stability assay. Representative results from two biologically independent experiments are presented. CHX, Cycloheximide. **(E)** GSEA plot showing enrichment of the Hallmark Hypoxia gene signature in the G367V upregulated genes determined by RNA-seq dataset. **(F)** Spectral flow cytometry analysis of CD4^+^Foxp3^+^ regulatory T (T_reg_) cells, with quantification of MFI for effector T_reg_ cell markers KLRG1, CTLA-4, GITR, and CD25 in MLL3 WT and G367V pre-onset tumors. MFI, Mean Fluorescence Intensity. N=14 tumors/group.

**Figure S2. Related to Figure 4.**
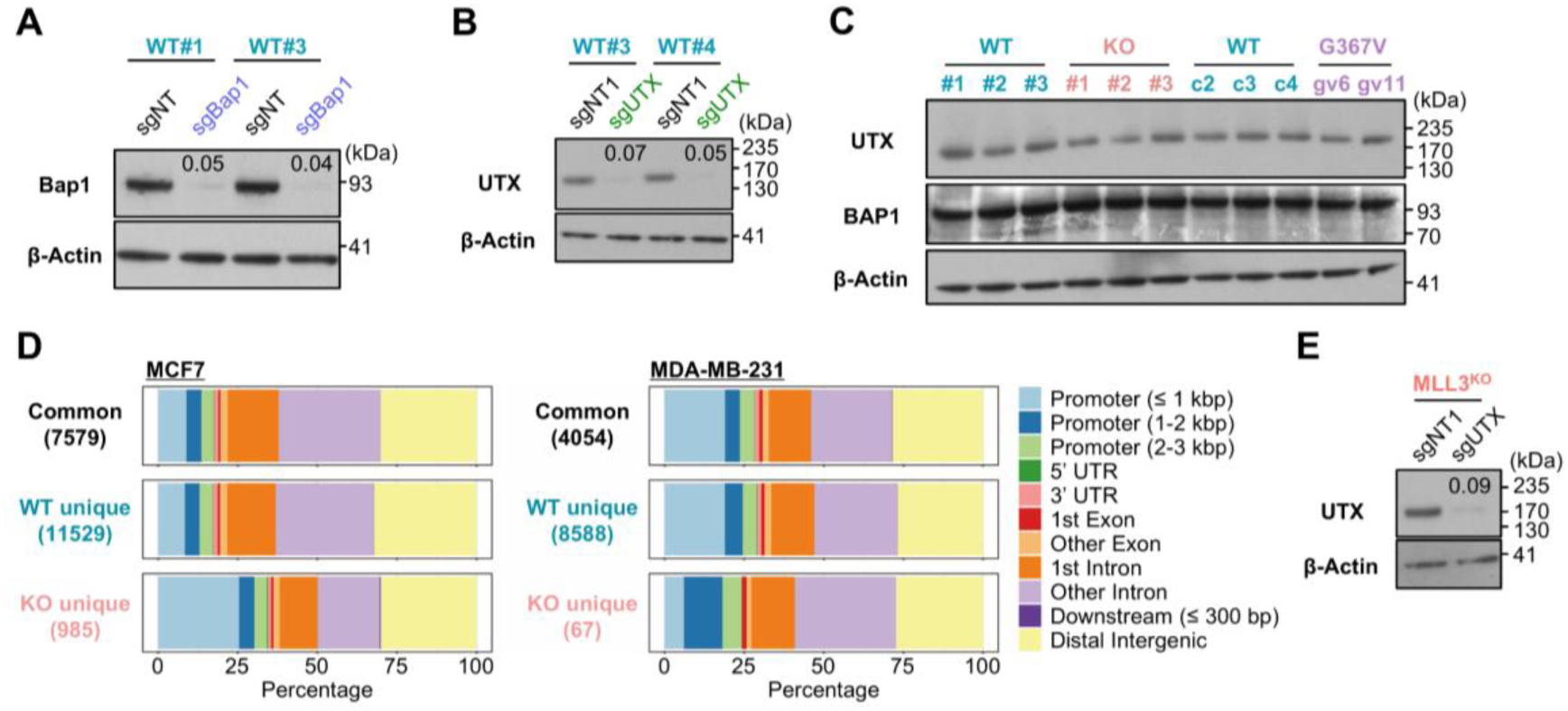
(A) WB analysis of BAP1 protein levels in sgNT or sgBap1 gRNA expressing MLL3 WT MaSCs. β-Actin was used as an internal control. The values show the relative intensity of BAP1 normalized to that of β-Actin against the sgNT group. **(B)** WB analysis of UTX levels in sgNT or sgUTX gRNA-expressing MLL3 WT MaSCs. β-Actin was used as an internal control. The values show the relative intensity of UTX normalized to that of β-Actin against the sgNT group. **(C)** WB analysis of total UTX and BAP1 protein levels in MLL3 WT, KO, and G367V mutant MaSCs. β-Actin was used as an internal control. **(D)** Genomic distribution of UTX binding sites (Common, MLL3 WT unique, and MLL3 KO unique peaks) in MCF7 and MDA-MB-231 human breast cancer cell lines, categorized by genomic features. **(E)** WB analysis of UTX levels in sgNT or sgUTX gRNA-expressing MLL3 KO MaSCs. β-Actin was used as an internal control. The values show relative intensity of UTX normalized to that of β-Actin against the sgNT cells.

**Figure S3. Related to Figure 5.**
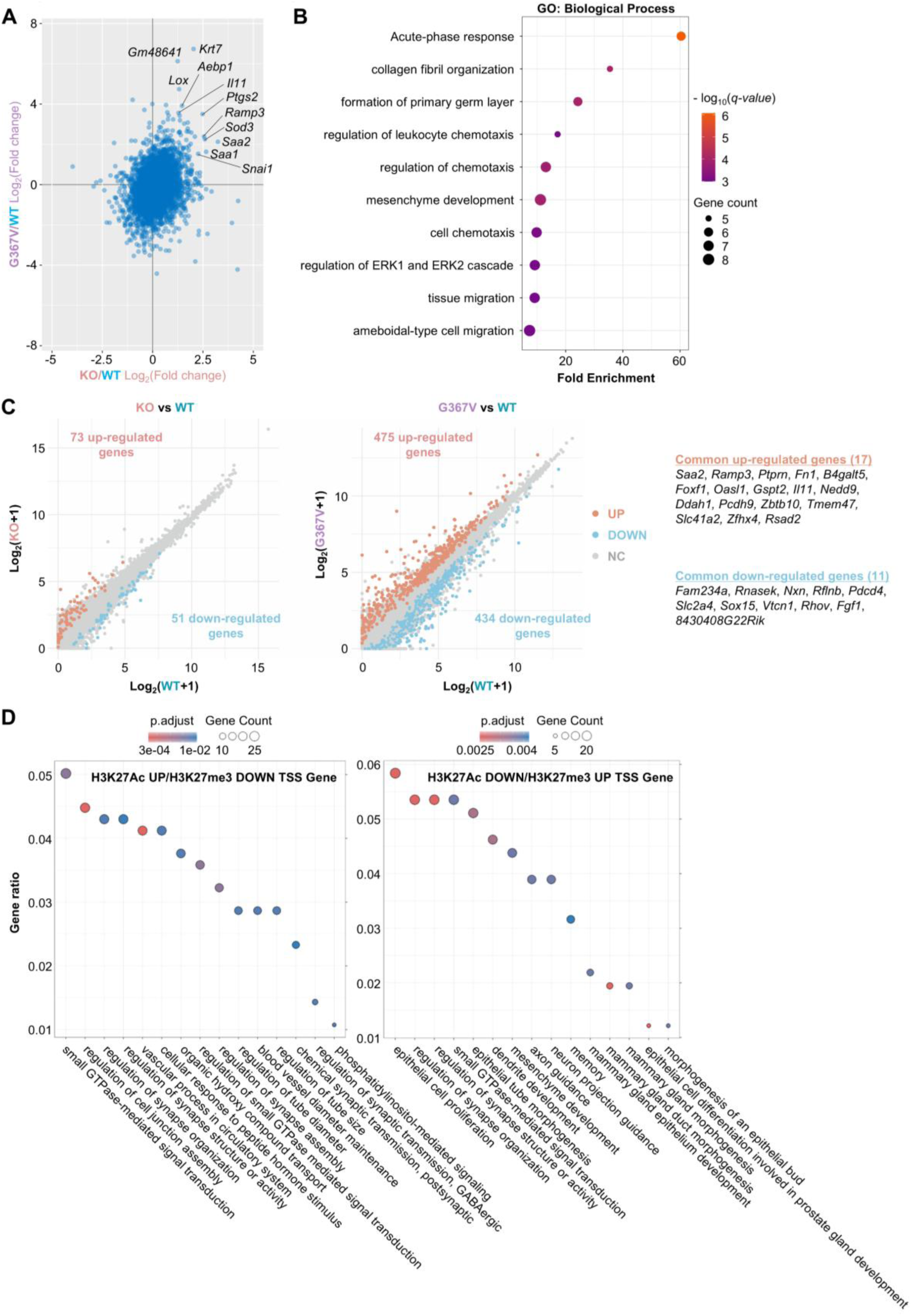
(A) Scatter plots comparing global gene expression changes in MLL3 KO and G367V MaSCs, relative to MLL3 WT MaSCs. **(B)** Gene Ontology (Biological Process) analysis of commonly up-regulated genes in both MLL3 KO and G367V MaSCs, highlighting terms such as acute-phase response and cell chemotaxis. **(C)** Differentially expressed genes between MLL3 WT and KO (left) or WT and G367V (right) MaSCs. Commonly up-regulated or down-regulated genes in both MLL3 KO and G367V MaSCs compared to MLL3 WT are listed on the right. **(D)** Gene ontology analysis with gene sets identified by CUT&TAG analysis that exhibited unique epigenetic status at around their promoter regions. GO Bioprocess terms were investigated.

